# Preventive Effects of Pomegranate Seed Oil on Transient Middle Cerebral Artery Occlusion via the Keap1/Nrf2/NQO1 Pathway in the rats Cortex

**DOI:** 10.1101/2024.09.13.612855

**Authors:** Dongyang Li, Chaoying Zhang, Qi Luo, Man Li, Meiqi Tian, Hanyi Jiao, Xiangwen Xi, Qifang Weng

**Affiliations:** Key Laboratory of Tropical Translational Medicine of Ministry of Education & Key Laboratory of Brain Science Research Transformation in Tropical Environment of Hainan Province, School of Basic Medicine and Life Sciences, Hainan Medical University, Haikou, 571199, China; The First Clinical College, Hainan Medical University, Haikou, 571199, China; School of Stomatology, Hainan Medical University, Haikou, 571199, China; Department of Cardiology, the Second Affiliated Hospital of Hainan Medical University, Haikou,570100, China

**Keywords:** transient middle cerebral artery occlusion, pomegranate seed oil, kelch like ECH associated protein 1, nuclear factor erythroid 2-related factor 2, NAD(P)H quinone oxidoreductase 1

## Abstract

Ischemic stroke remains a pressing challenge that needs to be solved. Energy metabolic failure is a critical factor contributing to mitochondrial dysfunction and oxidative stress in the pathogenesis of brain ischemia, leading to the generation of excessive reactive oxygen species. Pomegranate seed oil (PSO) exhibits antioxidant properties; however, its protective effects against cerebral ischemia/reperfusion injury and the underlying mechanisms remain unclear. In this study, a transient middle cerebral artery occlusion (tMCAO) rat model was employed to simulate cerebral ischemia/reperfusion injury. We investigated the mechanisms by which different concentrations of PSO modulate oxidative damage caused by cerebral ischemia/reperfusion injury through the Keap1/Nrf2/NQO1 pathway in cortex. SD male rats were randomly divided into four groups: Control, tMCAO+NaCl, tMCAO+LO (low concentration of PSO), tMCAO+MO (medium concentration of PSO), and tMCAO+HO (high concentration of PSO). Our findings suggest that low concentration of PSO exerts neuroprotective effects by activating Nrf2 and NQO1, thereby reducing oxidative stress. Furthermore, LO significantly improved neurological scores and reduced neuronal edema. Additionally, the results demonstrated an increase in superoxide dismutase (SOD) levels and a decrease in malondialdehyde (MDA) levels. In contrast, MO and HO exhibited suboptimal effects. To sum up, these results indicate that PSO activates neuroprotective pathways against oxidative stress following cerebral ischemia/reperfusion injury via the Keap1/Nrf2/NQO1 pathway, providing novel insights into potential preventive therapies for cerebral ischemia/reperfusion.

## Introduction

Stroke is the second leading cause of death and the third leading cause of death and disability in worldwide (1). Cerebral ischemia/reperfusion injury, which typically occurs during the treatment phase of ischemic diseases, is triggered by a blood reperfusion injury in the affected brain area. The complexities surrounding the etiology of cerebral ischemia/reperfusion are multifaceted, encompassing a range of risk factors including hypertension, atherosclerosis, and cardiac abnormalities, which contribute to the disruption of blood supply to the brain (2). Following cerebral ischemia, patients frequently suffer from edema and hemorrhagic transformation, exacerbating the injury and complicating clinical outcomes. The pathological processes involved in cerebral ischemia/reperfusion are regulated by intricate mechanisms that encompass excitotoxicity, inflammation (3), oxidative stress (4), and apoptosis (5). These interrelated factors play a critical role in the progression and severity of ischemic damage. Many experimental studies have shown that oxidative stress is one of the most important factors in cerebral ischemic reperfusion injury (6) (7). The NAD(P)H quinone oxidoreductase 1 (NQO1) activated by kelch like ECH associated protein 1/nuclear factor erythroid 2-related factor 2 (Keap1/Nrf2) pathway (8). The Keap1/Nrf2 pathway plays an important role in protecting cells against various stresses (9) (10). Nrf2 is an important redox-sensitive transcription factor. Under normal physiological conditions, Nrf2 is anchored in the cytoplasm by Keap1. However, Nrf2 dissociates from Keap1 and rapidly translocates into the nucleus during oxidative stress, then binding to the antioxidant response element to activate the transcription of antioxidant enzyme genes, thereby playing an antioxidative role (11). Recent research has shown that Keap1/Nrf2 pathway is one of the major cellular defense systems against oxidative stresses (12) (13).

Pomegranate seed oil (PSO) is obtained from pomegranate seeds, which contains polyunsaturated fatty acids, monounsaturated fatty acids, saturated fatty acids, and some bioactive compounds such as phenolic compounds, tocopherols and phytosterols. It has been reported that pomegranate and its derivatives as PSO have a protective effect via antioxidant, anti-inflammatory, and neuroprotective activities (14). The recent studies have reported that PSO reduces plasma interleukin-6 and tumors necrosis factor levels in high-fat-diet-induced obese mice (15). In addition, for multiple sclerosis, PSO improvement of cognitive dysfunction through the antioxidative effects both in patients (16) and mice (17). Furthermore, PSO nanoemulsion can significantly reduce neuronal death, alleviate the proliferation of glial cells, and prevent mitochondrial damage to prevent a cognitive and behavioral decline in traumatic brain injury mice (18). Punicic acid, one of main components of PSO, protects cardiac fibrosis via antioxidant activity by Nrf2 activation (19). Here in, we employed the tMCAO rat model to verify whether PSO effect by activating the Keap1/Nrf2/NQO1 pathway.

## Materials and Methods

All methodologies employed in this study were in accordance with the guidelines on the Use and Care of Laboratory Animals set by the National Institutes of Health and were approved by the Institutional Animal Care and Use Committees of Hainan Medical University.

## Experimental animals

Male Sprague-Dawley rats (6–8 weeks-old, 180–200 g) were used for this study, which were purchased from HCB Co., Ltd. (No. 85), (Changsha, China), (RRID: SCXK, 2022-0011). Animals were housed in room temperature of 21–23 °C and 12/12 h light/dark cycle with free access to food and water. All animal experiments were approved by the Ethics Committee of Hainan Medical University. After adaption for a week, the rats were randomly divided into sham operation and transient middle cerebral artery occlusion (tMCAO) groups (Fig. 1). Before tMCAO, the rats were given PSO twice a day for a total of 7 days, including 2.5 ml NaCl, low concentration of PSO (1.3 ml, LO), medium concentration of PSO (2.5 ml, MO), and high concentration of PSO (3.8 ml, HO). In preparing tMCAO, rats were anesthetized with chloral hydrate (300 mg/kg body weight, i.p). Positioned supinely with the head immobilized on a surgical platform, and the tongue was pulled out to one side. A midline incision in the neck was made to expose the bilateral common carotid arteries, and then, the bilateral common carotid arteries were ligated by vascular clamp for 20 min. The sham operation group received all the surgical procedures except for the occlusion of blood vessels. The surgical incisions were closed layer by layer with suture, after which the rats were returned to their individual cages with a 37 °C heating pad.

**Fig. 1.**
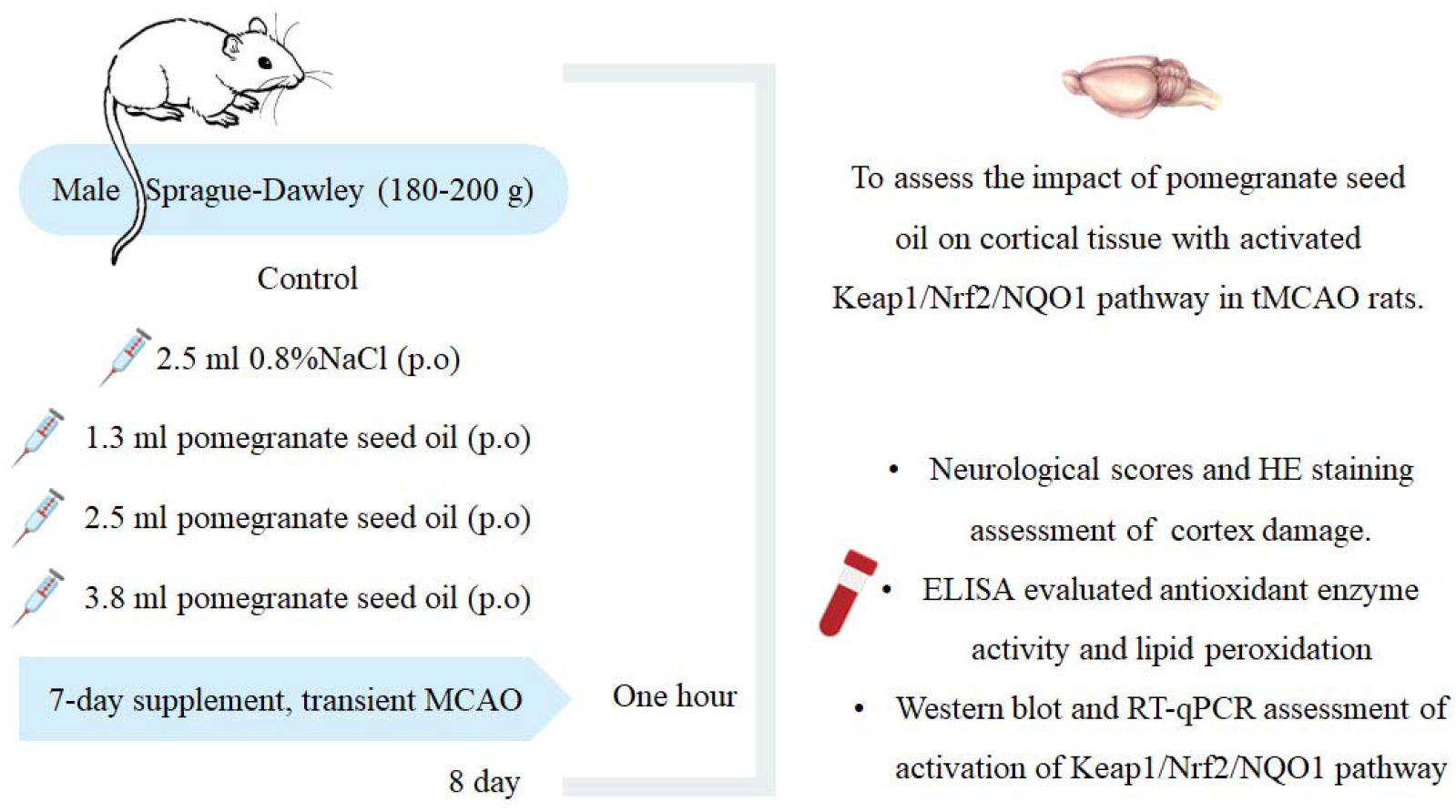
Flowchart of experiments.

## Evaluation of Neurological Injuries and Tissue Preparation

To evaluate brain damage, we conducted neurological assessments one hour prior to sampling. The neurologic examination was performed as follows: Normal-grade 0, no observable deficit; Moderate-grade 1, blepharoptosis; Severe-grade 2, paralysis, ataxia without blepharoptosis; Very severe-grade 3, blepharoptosis, paralysis and ataxia.

After one hour of tMCAO, rats were decapitated after anesthesia with chloral hydrate (300 mg/kg body weight, i.p), brains were quickly removed and soaked in ice-cold artificial cerebrospinal fluid that contained (in mM) 126 NaCl, 3 KCl, 1.3 MgSO_4_, 2.4 CaCl_2_, 1.3 NaH_2_PO_4_, 26 NaHCO3, 10 Glucose, 305 mEq/Kg, pH 7.4, and gassed with 95 % O_2_/5 % CO_2_. The cortex in one side of the brain was punched out to extract proteins for Western blots. The other side of the cortex was fixed in 4 % paraformaldehyde for 24 h for immunohistochemistry and Hematoxylin and Eosin Staining. For assaying plasma SOD and MDA levels, trunk blood was collected with heparinized test tubes. We did not observe any signs of peritonitis in rats after intraperitoneal injection of chloral hydrate.

## Hematoxylin and Eosin Staining

The tissues were dissected and fixed in 4 % paraformaldehyde and then used for immunohistochemistry and hematoxylin and eosin staining. After dehydration with graded alcohol, xylene was used to hyalinize the tissues before paraffin imbedding. Wax blocks were sectioned into 4-um thick sections by paraffin slicer (Leica, Wetzlar, Germany). For identifying the histological features of the cortex, the hematoxylin and eosin staining were the same as previously described (20). Following deparaffinization and rehydration, sections were stained with Hematoxylin solution for 5 min, treated with 1 % hydrochloric acid–alcohol for 3 s and stained with eosin solution for 30 s, dehydrated with graded alcohol solutions and cleared in xylene (C0105S, Beyotime, China). The sections were examined and photographed using a digital microscope (Slide Viewer, 3D histech).

## Western Blots

The methods were the same as previously described (20) (21). In brief, 35 μg of proteins were separated on 10% SDS-PAGE gels and transferred onto PVDF membranes. Protein membranes were pretreated with a protein free rapid blocking buffer (PS108) for 15 min at room temperature, and then incubated with antibodies against target proteins at 4 °C overnight. The primary antibodies included Nrf2 (PA5-88084, rabbit, 1:1000, Thermo Fischer Scientific), Keap1 (NBP1-83106, rabbit, 1:1000, novus), and NQO1 (NB200-209, mouse, 1:1000, novus). Loading controls were set using antibodies against β-actin (ab8226, rabbit, 1: 1000, abcam). The protein membranes were further incubated with HRP-conjugated secondary antibodies (AS038, Donkey Anti-Rabbit IgG, AS033, Donkey Anti-Mouse IgG, 1:10000, Abclonal). Protein bands were visualized with an automated chemiluminescence image analysis system (Tanon 5200, Shanghai).

## RT-qPCR Assay

Total RNA was extracted from rat brain tissue using TRIzol reagent (BS258A, Biosharp), with an incubation period of 5 minutes. Following this, the solution was centrifuged at a force of 12,000 ×g for 10 minutes. The supernatant, measuring 1.5 mL, was then amalgamated with 200 µL of chloroform in a centrifuge tube and further centrifuged at 12,000 ×g for another 10 minutes at a temperature of 4 °C. Subsequently, the supernatant was merged with 500 µL of isopropyl alcohol in a new centrifuge tube and centrifuged under the same conditions as before. After discarding the supernatant, the precipitate was cleansed with 1 mL of 75 % absolute ethanol, followed by another cleansing with 1 mL of absolute ethanol. The solution was then centrifuged at 7500 ×g for 5 minutes at 4 °C, after which the supernatant was removed, and the RNA was resuspended in 40 µL of DEPC water. The creation of cDNA was achieved by reverse transcription at 42 °C for a duration of 1 hour, followed by 75 °C for 5 minutes (K1622, RevertAid First Strand cDNA Synthesis Kit, Thermo). SYBR Green qPCR Master Mix (B21202, Bimake) was used for real-time fluorescence qPCR detection. The reaction system was assembled with 10 µL of UltraSYBR Mixture, 1 µL of PCR Forward Primer (10 µM), 1 µL of PCR Reverse Primer (10 µM), 1 µL of cDNA template and 7 µL of DEPC water. The RT-qPCR conditions were set as follows: a denaturation phase at 95 °C for 10 minutes, followed by 40 cycles of 15 seconds at 95 °C and 60 seconds at 60 °C. The primer sequences used for the RT-qPCR are presented in Table 1.

**Table 1.**
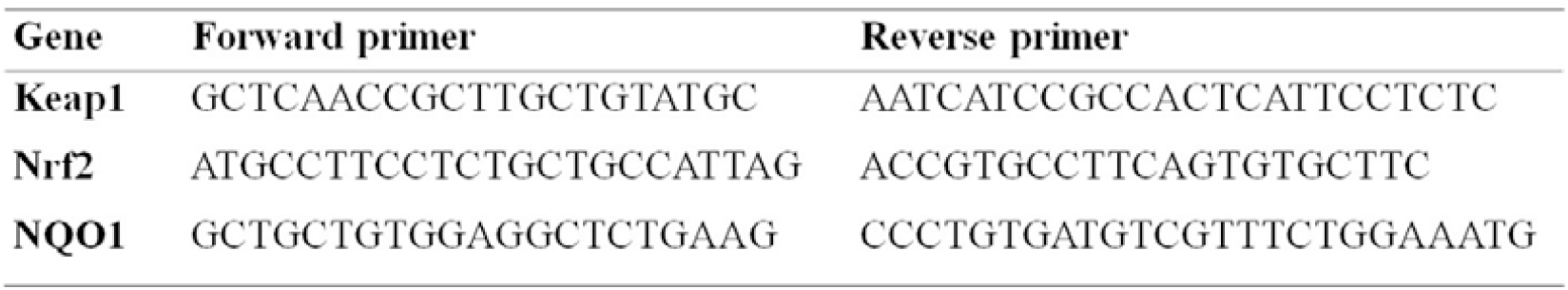
prime sequences required for RT-qPCR.

## Enzyme-Linked Immunosorbent Assay of SOD and MDA Levels

Assaying plasma superoxide dismutase (SOD, BC0170, Solarbio) and malondialdehyde (MDA, BC0025, Solarbio) levels used the method as instructed by the assay manual. Briefly, after adding standard or sample solutions (50 μl) to the assigned wells, 10 μl detection reagent and 40 ul sample diluent were added to all wells, and then add 100 μl of HRP-conjugate reagent to each well, cover with an adhesive strip and incubate for 60 min at 37 °C. After washing 5 times with 400 μl wash solution, 50 μl chromogen solution A and 50 μl chromogen solution B were added to each well to incubate for 15 min at 37 °C. The reaction was stopped by adding 50 μl stop solution. The optical density of the reaction was measured immediately at 450 nm. This assay kit can detect SOD levels as low as 1.0 pg/mL and MDA levels as low as 0.1 nmol/ml. It has no cross reaction with other soluble structural analogues.

## Data Analysis

Data analyses used the same methods as previous publication with minor modifications (22). Data were expressed as mean ± SEM using SigmaStat program 19 software (Chicago, IL), and *P <* 0.05 was considered statistically significant. ANOVA was used for comparison between multiple groups followed by Bonferroni test (for normally distributed variables) or Kruskal-Wallis test (intergroup comparison for not normally distributed variables) where appropriate.

## Results

### PSO ameliorates Neurological Scores and Cortical Damage after tMCAO

After verifying the success of the tMCAO model, the neurological index score was evaluated. The results showed that NaCl, LO and HO increased neurological scores compared with CTR rats (P < 0.01). However, the neurological scores of LO rats were lower (Fig. 2A). These results are consistent with the findings in both patients (23) and rat model (24).

**Fig. 2.**
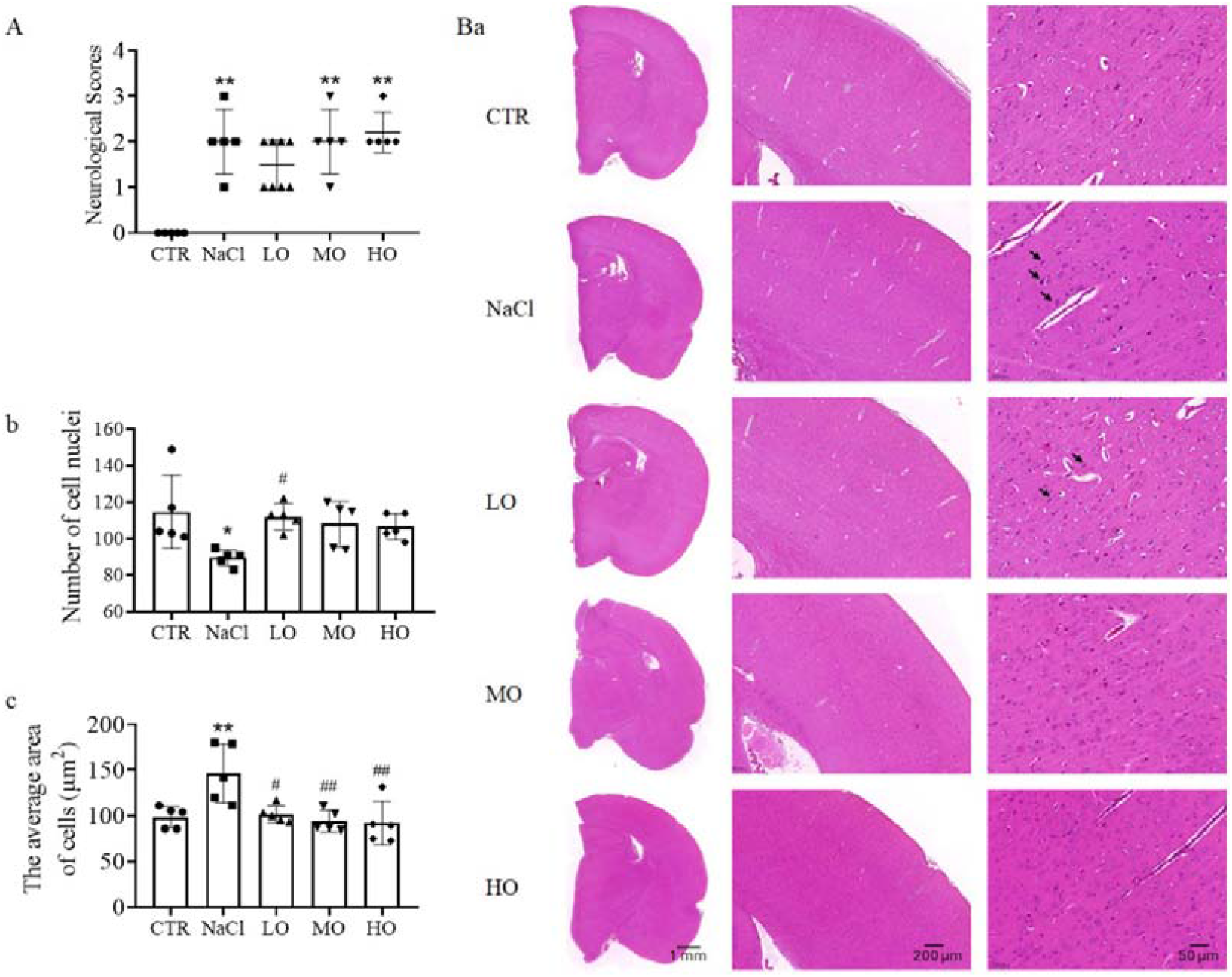
Effects of PSO on neurological scores and cortical damage after tMCAO. A. The score of neurologic examination grading in different groups. B. Hematoxylin– Eosin staining of the cortical (a) and the summary graph of statistical analysis of the number of cell nuclei (b) and the average areas of neurons (c). *, *P* < 0.05, **, *P* < 0.01, vs. control by One Way ANOVA. #, *P* < 0.05, ##, *P* < 0.01, vs. NaCl by One Way ANOVA. Abbreviations: CTR, control; LO, low concentration pomegranate seed oil; MO, medium concentration pomegranate seed oil; and HO, high concentration pomegranate seed oil.

Consistent with the changes in neurological scores, the cortical also changed can be caused by tMCAO (25). As shown in Fig. 2B, the nuclei underwent pyknosis in NaCl and LO groups after tMCAO. The number of nuclei significantly was reduced in NaCl group (P < 0.05). Moreover, there was significantly reversed in LO (P < 0.05). The number of nuclei were increased in MO and HO (P > 0.05). However, the retention of water in cells in NaCl group led to the expansion of cell volume, we found that the area of neurons was increased in NaCl group, and PSO improve neuronal edema (P < 0.01). As a whole, there was high consistent between the neurological scores and cortical damage.

### Evaluation of antioxidant enzyme activity and lipid peroxidation in PSO

SOD is a crucial antioxidant enzyme that protect cells from oxidative damage. MDA, a product of lipid peroxidation, which represents the degree of lipid peroxidation. SOD and MDA are important indicators for evaluating oxidative stress in terms of antioxidant capacity and oxidative capability, respectively (26). To understand the antioxidative effect of PSO, we evaluated the activity of SOD and MDA in plasma, That is, plasma SOD levels decreased significantly in NaCl, MO and HO groups (Fig. 3A). That is, SOD levels at NaCl were significantly lower than the CTR group (P < 0.05) and LO group can significantly reverse the change (P < 0.05). However, the increase of MO and HO group were not significant (P > 0.05). This suggests that PSO significantly alter plasma SOD levels in NaCl rats, indicating a potential beneficial effect of PSO on antioxidant enzyme activity in tMCAO conditions.

**Fig. 3.**
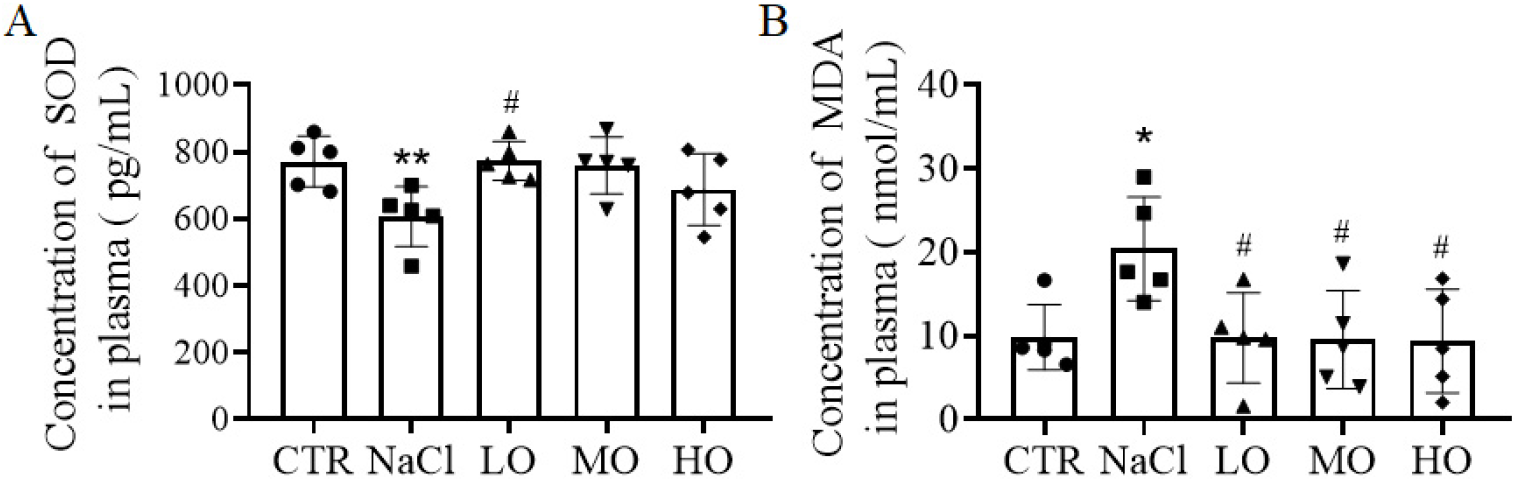
Plasma SOD and MDA levels after tMCAO. Effects of PSO treatment on SOD level (A) and MDA levels (B) in plasma. *, *P* < 0.05, vs. control by One Way ANOVA. #, *P* < 0.05, vs. NaCl by One Way ANOVA. Abbreviations: SOD, superoxide dismutase; MDA, malondialdehyde; CTR, control; LO, low concen00tration pomegranate seed oil; MO, medium concentration pomegranate seed oil; and HO, high concentration pomegranate seed oil.

There were significantly higher MDA levels in NaCl compared to the CTR (P < 0.05). However, MDA levels at LO, MO and HO (P < 0.05) were lower than those in the NaCl group (Fig. 3B). These results suggest that PSO treatment in tMCAO rats (NaCl) led to significant changes in plasma MDA levels compared with CTR rats, it results in a significant difference in plasma MDA levels between NaCl and PSO treated groups.

### The role of KEAP1/NRF2/NQO1 in tMCAO rats

Nrf2, a transcription factor, as one of the most important cellular defense mechanisms against oxidative stress. Keap1 is an endogenous inhibitor of Nrf2 also involved in oxidative stress (27). In theory, the protective effect of PSO can also be achieved by activities of Keap1/Nrf2 pathways. To test this hypothesis, we analyzed the protein and gene levels of components of the Keap1/Nrf2/NQO1 pathways in the cortex tissues of rats were evaluated by the western blotting and RT-qPCR. The Keap1 protein levels in NaCl, MO and HO groups were significantly increased (P < 0.05).

However, LO group shows a decreasing trend (P > 0.05, Fig. 4A). Moreover, Nrf2 was lowest in NaCl rats (P < 0.05), the PSO shows a trend of reversing Nrf2 protein levels (P > 0.05, Fig. 4C). Furthermore, NQO1 protein also expressed lowest in NaCl group (P < 0.05) and significantly increased in LO group (P < 0.05), the MO and HO groups was no difference (P > 0.05, Fig. 4E).

Further analyses the gene expression of Keap1, Nrf2 and NQO1 in the cortex tissues of rats, the result is consistent with the protein expressions. That is, Keap1 mRNA significantly increased in NaCl group and reduced in LO group (P < 0.05); however, the expression of Keap1 mRNA in HO group is different to the expression of protein (Fig. 4B). In addition, the Nrf2 mRNA levels were significantly lower in NaCl, MO and HO groups compared with the CTR group (P < 0.05); LO group reversed the expression of Nrf2 (P > 0.05, Fig. 4D). Similar to the findings in the expression of NQO1 protein, NQO1 mRNA levels were significantly decreased in the cortex tissues of rats from the NaCl group compared with the CTR group (P < 0.05); The NQO1 mRNA levels were significantly increased in the PSO that in low concentration (P < 0.05), not medium and high concentration (P > 0.05, Fig. 4F). Taken together, these results are consistent with the hypothesis that PSO in low concentration attenuates tMCAO by NQO1 via activation of Nrf2.

**Fig. 4.**
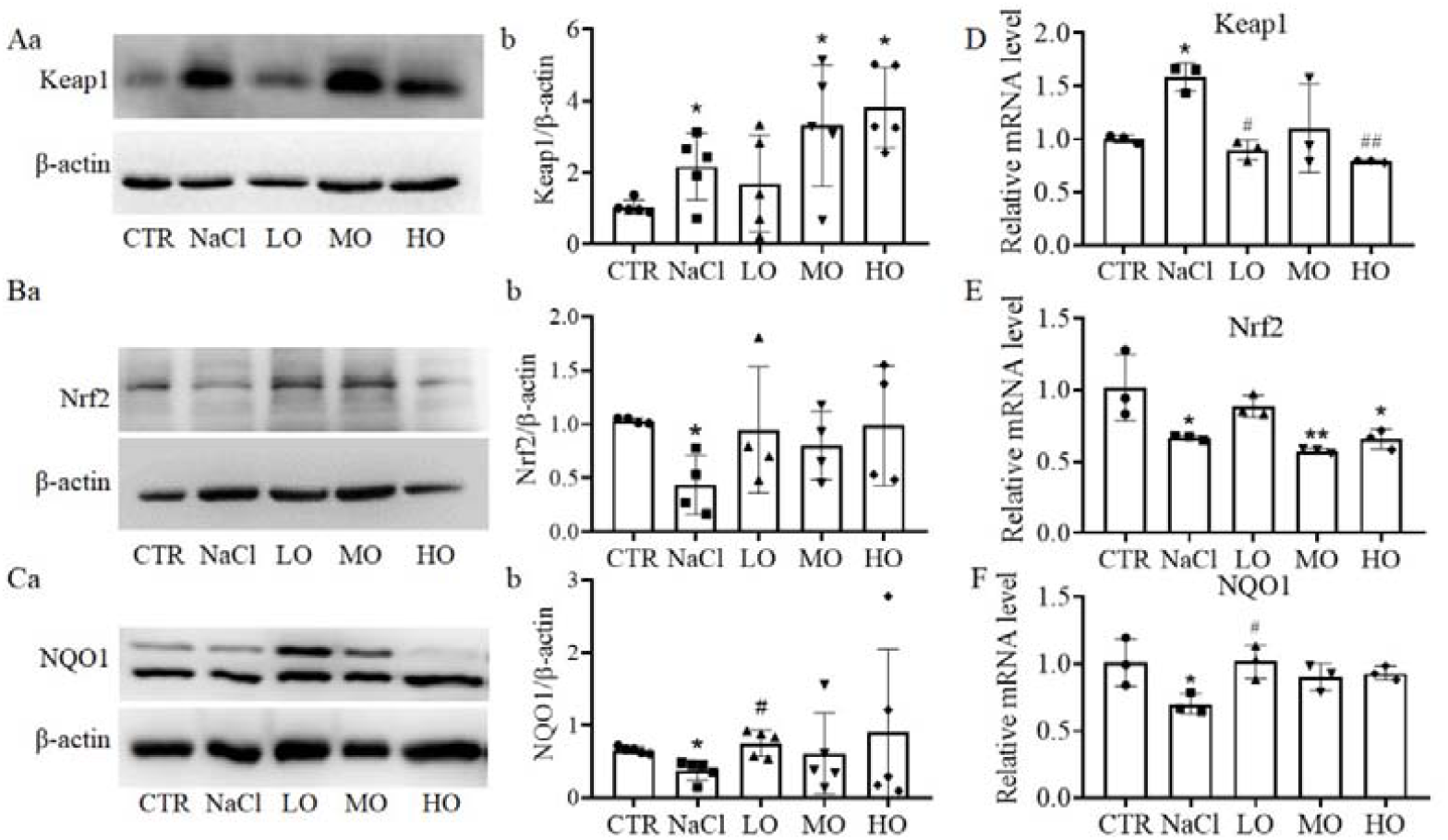
Effects of PSO on Keap1-Nrf2-NQO1 pathway after tMCAO. A-C. Western blotting bands (a) and the summary graphs (b) showing the effects of PSO treatment on Keap1 (A), Nrf2 (B) and NQO1 (C) protein levels. D-F. Keap1 (D), Nrf2 (E) and NQO1 (F) in cortex. *, *P* < 0.05, **, *P* < 0.01, vs. control by One Way ANOVA. #, *P* < 0.05, ##, *P* < 0.01, vs. NaCl by One Way ANOVA. Abbreviations: Keap1, kelch like ECH associated protein 1; Nrf2, nuclear factor erythroid 2-related factor 2; NQO1, NAD(P)H quinone oxidoreductase 1; CTR, control; LO, low concentration pomegranate seed oil; MO, medium concentration pomegranate seed oil; and HO, high concentration pomegranate seed oil.

## Discussion

In this study, we utilized tMCAO rat model to investigate the mechanisms underlying cerebral ischemia/reperfusion injury. This model effectively simulates the pathological processes involved in ischemic strokes, providing a foundation for exploring potential therapeutic interventions. Our research particularly focused on the correlation between tMCAO and oxidative stress responses, revealing significant oxidative stress following ischemic injury. Notably, our findings indicate that PSO exhibits strong antioxidant properties, effectively mitigating the oxidative damage associated with cerebral ischemia/reperfusion. Furthermore, we explored the mechanisms through which PSO exerts its protective effects, specifically examining its role in the Keap1/Nrf2/NQO1 pathway. The results highlight that PSO can enhance the antioxidant response through Keap1/Nrf2/NQO1 pathway, thereby improving the brain’s resilience against oxidative injury. These results of this study provide valuable insights into potential preventive strategies for mitigating damage from tMCAO.

tMCAO is a widely utilized experimental model to study the mechanisms of cerebral ischemic injury and its consequence (28). Following ischemia and subsequent reperfusion, significant neuronal damage occurs, primarily leading to devastating clinical symptoms such as hemiparesis, weakness in limbs, and slurred speech in patients (23). The neurological deficits focus on the severe impact of ischemic injury on brain tissue and functional outcomes. In our study, we observed behavioral abnormalities in rats of tMCAO (Fig. 2), accompanied by notable cortical damage.

Importantly, the administration of low concentration of PSO improve adverse effects associated with tMCAO, positively influencing both the extent of brain injury and behavioral alterations. Conversely, medium and high concentration of PSO did not exhibit significant improvements, suggesting a dose-dependent relationship in its therapeutic effects. The promising results associated with low concentration of PSO suggest its potential as a pharmacological agent for the prevention of ischemic damage. However, these findings require further clinical investigations to confirm its efficacy and safety in human. Continued exploration in this area could provide essential insights and contribute to developing novel therapeutic strategies for cerebral ischemia/reperfusion patients.

One of the primary mechanisms underlying cerebral ischemia/reperfusion injury is oxidative stress, which leads to a cascade of pathological processes to neuron. The abrupt restoration of blood flow after a period of ischemia results in the excessive generation of reactive oxygen species (29), diminishing the antioxidant defenses and resulting in cellular damage. This oxidative stress is intimately linked to neuronal dysfunction and death. PSO has been recognized for its antioxidant properties, showing efficacy in preventing various conditions, including neurodegenerative diseases (30), colitis (31), and diabetes (32). Its potential to mitigate oxidative stress in the cerebral ischemia/reperfusion injury is worth exploring. In our study, we found that following cerebral ischemia/reperfusion, levels of SOD, a crucial antioxidant enzyme, were significantly reduced. This is consistent with previous study (33).

Notably, the administration of low concentration of PSO can improve this decline in SOD activity, suggesting its potential role in enhancing the brain’s antioxidant capacity. Conversely, medium and high concentration of PSO didn’t demonstrate any significant changes in SOD levels, indicating a possible threshold effect on the dosage. Additionally, we observed a significantly increase in MDA, a byproduct of lipid peroxidation, following tMCAO. Interestingly, rats in the LO, MO, and HO groups all showed efficacy in reducing MDA levels, suggesting a broad capacity for PSO to alleviate oxidative stress. However, the mechanisms that PSO elevates oxidative damage in the cerebral ischemia/reperfusion remain to be elucidated. Further studies on the specific bioactive compounds in PSO and their interactions with oxidative stress pathways are necessary to fully understand this potential therapeutic effect.

The Keap1/Nrf2/NQO1 pathway is a vital regulatory system in managing oxidative stress (13) (27) and has been implicated in various neurological disorders, including cerebral ischemia/reperfusion injury (9), epilepsy (10), and neurodegenerative diseases such as Alzheimer’s disease, Parkinson’s disease, and multiple sclerosis (34) (35), as well as subarachnoid hemorrhage (36). Activation of this pathway helps to upregulate antioxidant proteins, thereby providing cellular protection against oxidative damage. In our findings, we observed a significant increase in both the protein and mRNA levels of Keap1 in the cortex following tMCAO. The administration of low concentration of PSO effectively mitigated these changes, illustrating its potential to modulate the Keap1/Nrf2 pathway. Conversely, medium and high concentration of PSO did not produce any significant changes in Keap1 levels. Additionally, we found that the proteins and mRNA expressions of Nrf2 and NQO1 were significant reduced after tMCAO, which were significantly improved with low concentration of PSO treatment, while medium and high doses also failed to show beneficial effects. These results suggest that PSO can enhance the antioxidant response through the Keap1/Nrf2/NQO1 pathway, potentially reducing oxidative stress associated with cerebral ischemia/reperfusion injury.

This mechanism highlights the promising clinical significance of low concentration of PSO as a preventive therapeutic agent against oxidative damage from ischemic damage, warranting further clinical investigations to explore its efficacy in humans.

## Conclusions

As a whole, tMCAO causes brain damage primarily through oxidative stress. Low concentration of PSO effectively enhance antioxidant responses via the Keap1/Nrf2/NQO1 pathway, while medium and high doses show minimal improvement. This study offers unique insights for clinical preventive strategies against cerebral ischemia/reperfusion injury.

